# Integration of a radiofrequency coil and commercial field camera for ultra-high-field MRI

**DOI:** 10.1101/2021.09.27.462001

**Authors:** Kyle M. Gilbert, Paul Dubovan, Joseph S. Gati, Ravi S. Menon, Corey A. Baron

## Abstract

**Purpose:** To develop an RF coil with an integrated commercial field camera for ultra-high field (7 T) neuroimaging. The RF coil will operate within a head-only gradient coil and be subject to the corresponding design constraints. The RF coil can thereafter be used for subject-specific correction of k-space trajectories—notably in gradient-sensitive sequences such as single-shot spiral imaging.

**Methods:** The transmit and receive performance was evaluated before and after the integration of field probes, while field probes were evaluated when in an optimal configuration external to the coil and after their integration. Diffusion-weighted EPI and single-shot spiral acquisitions were employed to evaluate the efficacy of correcting higher order field perturbations and the consequent effect on image quality.

**Results:** Field probes had a negligible effect on RF-coil performance, including the transmit efficiency, transmit uniformity, and mean SNR over the brain. Modest reductions in field-probe signal lifetimes were observed, caused primarily by non-idealities in the gradient and shim fields of the head-only gradient coil at the probe positions. The field monitoring system could correct up to second-order field perturbations in single-shot spiral imaging.

**Conclusion:** The integrated RF coil and field camera was capable of concurrent field monitoring within a 7T head-only scanner and facilitated the subsequent correction of k-space trajectories during spiral imaging.

## 1 Introduction

MRI relies on the integrity of static and dynamic gradient fields to accurately encode and subsequently reconstruct images using Fourier acquisition schemes. Imperfections in the gradient field can be caused by the MR system and the subject, on both short and long time scales^1^. System imperfections include gradient amplifier propagation delays, gradient eddy currents^2,3^, heating (*B*_0_ drift)^4^, and mechanical vibration^5^, while patient-induced field perturbations are typically induced by motion^6^, whether respiratory or sporadic. These imperfections in the gradient field can result in image artifacts, such as blurring, ghosting, and distortion; however, with the correction of the encoding trajectories, these artifacts can be mitigated^7–9^.

Field cameras^1,10^ have been developed for such a purpose: they monitor the spatially and temporally varying magnetic field, using this information to retrospectively correct perturbations in k-space when reconstructing images^11^. Field cameras often rely on a set of small NMR coils (field probes^10,12–17^) strategically placed about the region of interest^18^. The phase evolution of a field probe’s free-induction decay (FID) provides a temporally dependent measurement of the local magnetic field. By sampling the magnetic field from multiple probes simultaneously, the magnetic field can be formulated as a series of weighted spherical harmonics to estimate its full spatial dependence^19^. To this end, the number and appropriate placement of probes will dictate the efficacy of measuring higher order field perturbations^20^.

Field monitoring can be performed either sequentially or concurrently. In the former, field cameras characterize gradient fields in the empty scanner^18^, with the measured field dynamics used to correct images from a subsequent acquisition as long as the acquisition parameters are the same^21,22^; however, this approach does not permit monitoring of patient-induced field perturbations or changes in scanning protocols (such as orientation, resolution, etc.) and requires a separate calibration acquisition—a workflow that is not technologist friendly. Although gradient impulse response functions (GIRFs) can be utilized to preclude the need for a separate scan^21^, they are limited to the correction of linearly time-invariant perturbations that may not be realized in practice^23^. Field probes have therefore been incorporated into external positioning devices and implemented in conjunction with existing coils^24,25^ for concurrent field monitoring. As a natural progression to this concept, 3T RF receive coils have been developed with probes integrated within the coil housing^26,27^. This architecture improves user workflow, while also allowing the coil to be designed with consideration for probe positioning, thereby optimizing performance of both the RF coil and the field probes. Concurrent field monitoring has been demonstrated to produce improvements in functional^28,29^ and anatomical^30^ imaging, with spiral encoding being particularly well-served by this correction technique due to its sensitivity to non-idealities in the gradient trajectory^29^. Diffusion-weighted imaging^31,32^ can achieve substantial SNR gains by leveraging spiral encoding^33^, making it an ideal application of concurrent field monitoring^34^.

Diffusion-weighted imaging has benefited from recent technological advances^35^, as it relies on high SNR and gradient strengths (b-values) to probe differences in molecular motion caused by the local anatomy. These technical advances include the adoption of ultra-high-field imaging to improve SNR/resolution and an improvement in gradient technology, including dedicated head coils^36^, to provide gains in gradient strength and slew rate—the confluence of which has resulted in higher b-values and shorter echo times, as demonstrated at 3T^37,38^ and 7T^39^. These advancements (ultra-high field strengths and head-only gradient coils) nonetheless create commensurate challenges to integrating a field camera within an RF coil: namely, (i) the ideal region of the gradient and shim fields, wherein field probes should ideally be located, is reduced, and (ii) the physical space to incorporate all the constituent components—i.e., the local transmit coil (a common requisite at ultra-high field), receive array, and field probes—is reduced, which can lead to increased coupling.

This manuscript describes a coil solution that integrates the field probes from a commercial field-monitoring system into a transmit/receive RF coil designed for operation within a commercial 7T head-only scanner. The coil topology is described with reference to the challenges imposed by its implementation within a head-only gradient coil at ultra-high field. As such, the transmit and receive performance is evaluated before and after the integration of the field probes, while the performance of the field probes is evaluated when in an optimal configuration external to the coil and after integration into the RF coil. The efficacy of correcting higher order field perturbations, and the resultant effect on image quality, is evaluated using diffusion-weighted EPI and single-shot spiral acquisitions.

## 2 Methods

### 2.1 Coil design

A holistic design approach was adopted to maximize the performance of the three subcomponents of the RF coil—the transmit coil, receive coil, and field probes—with deference to the limitations imposed by the head-only gradient coil. The coil was also designed to be practical for routine use by minimizing additional setup time required by the operator.

#### 2.1.1 Coil housing

The coil housing—identical to that previously described^40^—was designed to create an unobstructed visual field with a 14.5-cm-wide gap above the eyes (Figure 1). The housing was split into anterior and posterior halves to facilitate the positioning of subjects. The anterior and posterior halves were overlapped to maximize geometric decoupling of receive elements^41,42^. Electrical connections between the two halves, for both transmit and receive coils, were made through built-in high-density connectors (ODU-MAC product line). The computer-aided-design file of the RF coil has been made publicly available (https://doi.org/10.17605/OSF.IO/7TSJM).

**Figure 1.**
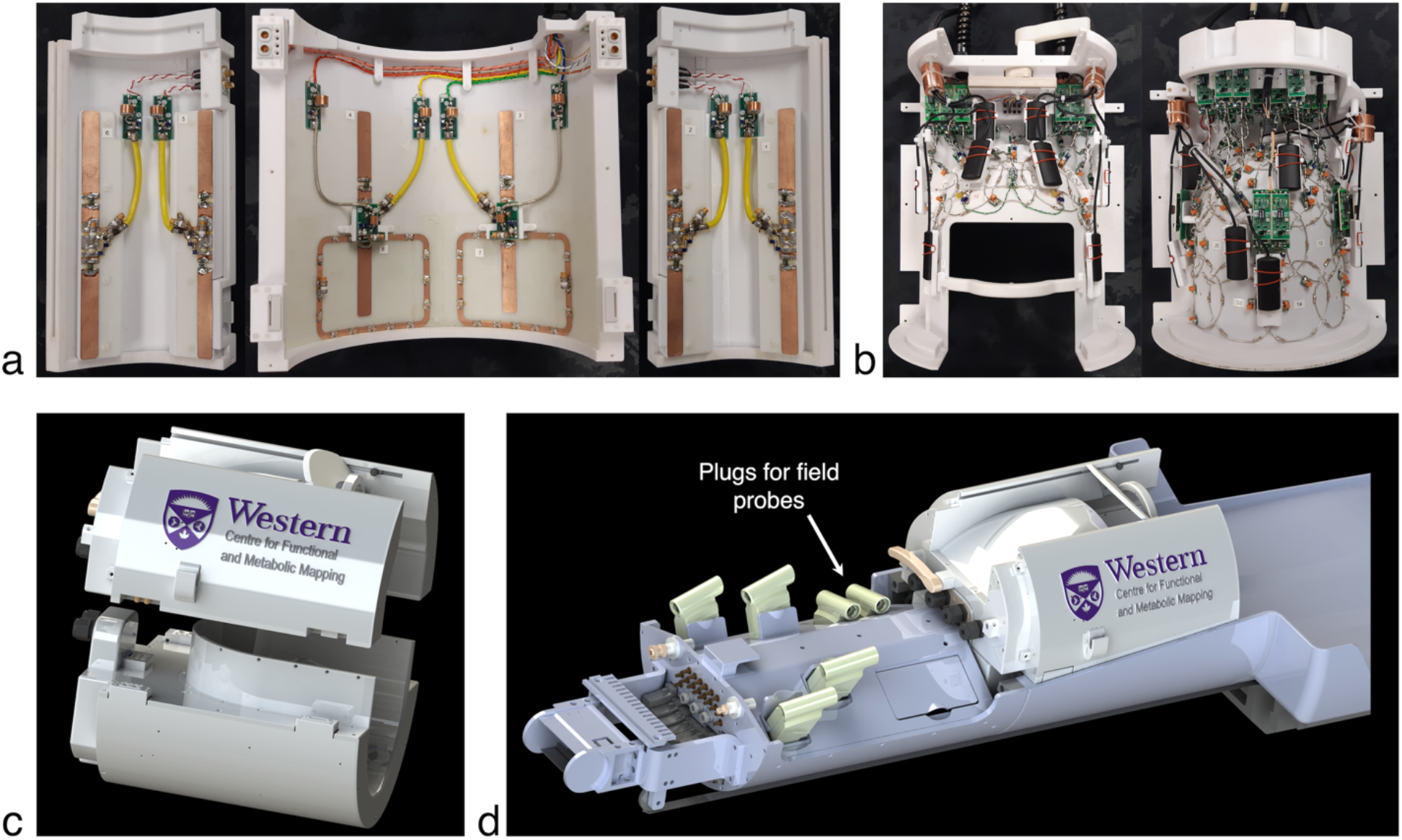
(a) The transmit coil is comprised of eight elements: six inductively shortened dipoles distributed about the circumference and two self-decoupled loops posterior to the cerebellum. (b) The 32-channel receive coil has field probes mounted directly to its housing in optimized locations. (c) The housing is a two-piece design to facilitate subject positioning, with electrical connections made through high-density connectors. (d) A custom socket was integrated into the scanner’s parallel-transmit interface to accommodate the two additional field-probe plugs.

#### 2.1.2 Transmit coil

The transmit coil was comprised of 8 elements: 6 dipoles (1.6-cm wide), equally spaced about the circumference of the housing, and 2 loops (13.1-cm wide by 10.2-cm high; 0.64-cm-wide traces) located posterior to the cerebellum. The transmit-coil topology was chosen for two reasons: (i) the flexibility in placing longitudinal dipoles allowed for a substantial gap in front of the eyes to create an unobstructed line-of-sight, and (ii) the posterior loops added a differential in sensitivity along the superior-inferior axis, which has been shown to improve *B*_1_^+^shimming capabilities over the whole brain^43,44^.

The anterior four dipoles were located sufficiently close to the head to produce consistent reflection measurements between subjects, while maintaining adequate distance to the receive coil and field probes to avoid high degrees of coupling. The posterior dipoles and loops were placed on the inside of the housing cover to increase the distance between the transmit and receive coils, as required for placing the field probes. The distance between the transmit and receive coils ranged from 2.3 – 3.8 cm; the distance between the transmit elements and isocentre ranged from 13.6 – 15.8 cm.

Posterior dipoles were physically shortened by 1.9 cm at their inferior aspect to minimize coupling between the high conservative electric fields at their ends to the overlapping loops—all other dipoles were 21.6-cm long. Each half-wavelength dipole was electrically shortened with wire-wound inductors to resonate at 297.2 MHz. A parallel variable capacitor (1 – 25 pF; Johanson 59H01) allowed for impedance matching to 50 Ω. An electrically shortened bazooka balun and a distal choke balun were attached to each transmit element to prevent common-mode currents. Active detuning was accomplished with a serial, high-power PIN diode (Chelton DH80106-44N) forward biased through RF chokes during transmission.

The virtual ground of each loop was aligned with the overlapping dipole to minimize their mutual coupling. The self-decoupling technique^45^ was implemented in loops to minimize their coupling with neighbouring elements, with capacitor values determined through simulation to minimize coupling with adjacent elements. Common-mode currents were mitigated by inclusion of a choke balun at the drive point. Active detuning was accomplished by identical means as for dipole elements. Circuit schematics of a transmit dipole and loop are provided in Figures 2a, b.

**Figure 2.**
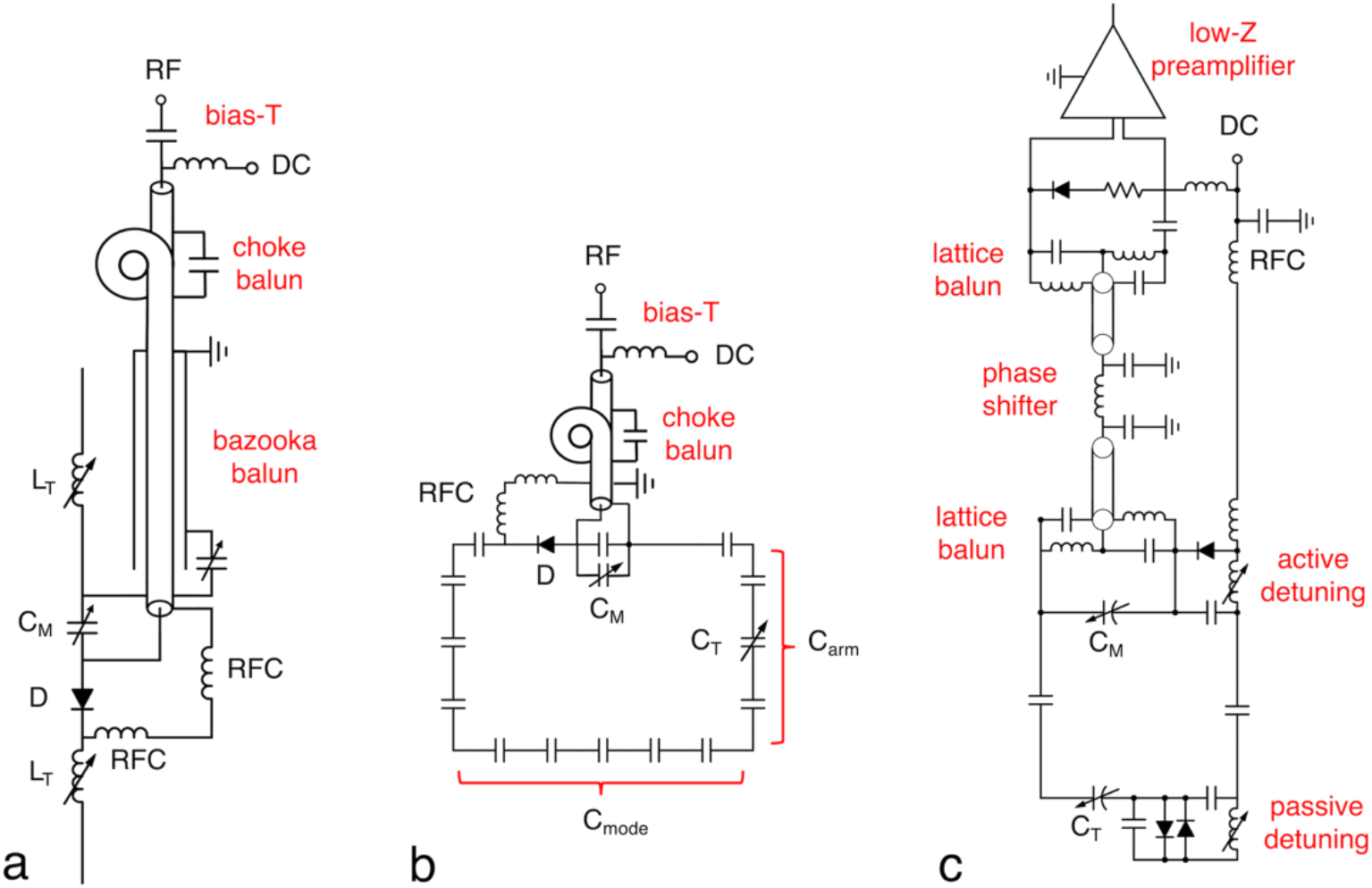
Circuit schematics of a (a) transmit dipole, (b) transmit loop, and (c) receive loop. Transmit dipoles were inductively shortened and included a serial active-detuning network. Transmit loops employed the self-decoupling technique, with C_arm_ and C_mode_ chosen to minimize coupling to adjacent elements. Receive loops employed both active and passive detuning and were connected to low-input-impedance preamplifiers by half-lambda coaxial cables. C_M_, matching capacitor, C_T_: tuning capacitor, D: diode, DC: direct current, L_T_: tuning inductor, RFC: radiofrequency choke.

The creation and validation of local SAR models followed the procedure described previously by the authors^40^. In brief, full-wave electromagnetic simulations of the transmit coil were performed when loaded with a phantom of known geometry, conductivity, and permittivity. Simulated and experimental *B*_1_^+^ maps were subsequently compared to validate the simulation model. With the accuracy of the simulation validated, the SAR of a body model (CST’s Hugo) was calculated to create local SAR matrices (*Q*-matrices), which were subsequently compressed into virtual observation points^46^ (VOPs) and used for online SAR monitoring. A safety factor of 1.8 was included in the VOPs to provide a conservative estimate of local SAR.

#### 2.1.3 Receive coil

The receive coil was comprised of 32 loop elements, arranged in a soccer-ball geometry^47^, with adjacent elements overlapped to reduce inductive coupling^41^. Elements were tuned to 297.2 MHz and matched to 75 Ω (i.e., the optimal impedance to minimize the noise figure of preamplifiers). A parallel-resonant, active-detuning circuit with a high-power PIN diode (Macom MA4P7435NM-1091T) was placed near the input of each receive element. A second active-detuning circuit was incorporated into the element encircling the eyes due to its large size. Parallel-resonant, passive-detuning circuits^48^ were added to each element as a secondary detuning method and to enhance detuning during transmission. Passive and active detuning networks were placed away from transmit dipoles and field probes, where possible, to reduce coupled power in regions of high electric field. A circuit schematic of a receive element is provided in Figure 2c.

Most preamplifiers were located at the superior aspect of the coil to conserve the already limited space for the integration of field probes. Elements were connected to these preamplifiers through half-lambda cables, including two lattice baluns and a phase shifter. Preamplifiers (Stark Contrast, Erlangen, Germany) had a low input impedance for preamplifier decoupling^41^ and an integrated cable trap between the first and second amplification stage. The inferior row of elements on the posterior half of the coil had preamplifiers placed directly at their output: this prevented excessively long cables, with non-amplified signal, within the transmit field. Preamplifiers were located away from transmit elements to avoid performance loss. Receive cables were routed between dipoles to minimize coupling to the transmit elements and, where possible, avoid proximity to field-probe cables.

#### 2.1.4 Field probes

A commercial field camera (“Clip-On Camera”; Skope, Zurich, Switzerland) was integrated into the RF coil. The field camera consisted of 16 ^19^F transmit/receive NMR probes attached to a dedicated spectrometer.

Field-probe locations were iteratively determined, with special consideration for the physical constraints of the RF coil and the limited imaging region of the head-only gradient coil (22-cm diameter sphere volume (DSV)). Initially, the ideal probe arrangement for measuring the spherical-harmonic representation of field perturbations (akin to being on the manufacturer’s positioning scaffold) was scaled to a radius that would allow the placement of field probes several millimetres beyond the inner housing of the receive coil. This minimized their distance to isocentre while avoiding being “too close” to materials that could cause local field inhomogeneities by virtue of their magnetic susceptibility. Field-probe locations were then modified to avoid placing them directly beneath transmit elements, near on-coil receive preamplifiers, or close to conductors. The inferior 3 field probes were 2 – 3 cm beyond the linear region of the head-only gradient coil in this configuration; therefore, they were translated 3 – 4.5 cm in the superior direction. The mean distance between field probes and isocentre was 13.5 cm (range: 11.9 – 14.9 cm)—i.e., outside the ideal linear region of the gradient coil, but unavoidable given the requisite size of a head RF coil. (Positions of the field probes are included in the publicly available CAD file.) After integration of field probes, both the transmit and receive coils were retuned and matched to compensate for minor changes.

Cabling of field probes to the field-camera spectrometer was particularly challenging^49^ due to the space-constraints within a head-only gradient coil and the desire to make the coil user-friendly to promote routine use. The parallel-transmit interface, located directly behind the bed’s head cradle, prevented the field-probe preamplifiers from being placed near the coil without occluding the subject’s line-of-sight during rear projection of visual stimuli. Field-probe preamplifiers were therefore placed outside of the magnet bore and underneath the false floor at the service end of the scanner. Field-probe cables within the coil were bundled into sets of four, with each bundle passing through a floating sleeve balun^50^ to suppress common-mode currents. The anterior and posterior cables were each terminated with an eight-channel ODU plug that was connected to a custom socket integrated into the parallel-transmit interface (see Figure 1)—this meant the operator was only required to connect two additional high-density plugs to utilize the field camera. Cables were routed through the parallel-transmit interface, secured to the top of the energy chain, then connected to the field-probe manufacturer’s shielded preamplifier box.

### 2.2 Imaging

All imaging was performed at the Centre for Functional and Metabolic Mapping at the University of Western Ontario. MR data collection was performed on a human, head-only 7T magnet with a scanner console operating on the VE12U software platform (Siemens Healthineers AG, Erlangen, Germany). The system is equipped with 32 receivers, 8 × 2-kW RF power amplifiers, and an AC84II head gradient coil (maximum gradient strength: 80 mT/m, maximum slew rate: 400 (T/m)/s; DSV: 22 cm) with a 36-cm-diameter clear bore (Siemens Healthineers AG, Erlangen, Germany). The gradient coil included a slotted RF shield built into its 40-cm inner diameter.

All human imaging was performed in accordance with the human research ethics board of the University of Western Ontario. Rigorous testing of both coil and patient safety was performed before the scanning of human subjects to ensure robust operation and subject safety during everyday use. Exhaustive testing followed regulatory guidelines specified by the IEC^51,52^, summarized by Hoffmann et al.^53^, and described in a previous study by the authors^40^.

### 2.3 Performance evaluation

All coil-performance assessments listed below were measured before and after integration of the field probes. Performance metrics were recorded for both a head and a head-mimicking phantom; however, since performance metrics were similar in both instances, only head data is presented.

#### 2.3.1 Transmit coil

Active detuning of the transmit coil was measured on the bench by driving an individual element from the transmit socket and measuring the transmitted signal with a pickup loop when the element was tuned and detuned. Scattering parameters were recorded from the directional couplers in the transmit path of the scanner when the coil was loaded with a human subject. Transmit efficiency maps of individual transmitters were subsequently acquired with the actual flip-angle imaging method^54,55^: matrix size: 96 × 96 × 72, FOV: 240 × 240 × 216 mm, TE/TR1/TR2: 1.7/20/100 ms, flip angle: 70°, BW: 1,410 kHz/pixel.

#### 2.3.2 Receive coil

The receive coil was tuned and matched when loaded with a head-mimicking phantom (ε_r_: 53, σ: 0.63 S/m; recipe derived from Duan et al.^56^) and placed inside a copy of the gradient coil’s RF shield. Preamplifier decoupling and active detuning of each receive element was measured using standard double-probe techniques^57^. The coupling, *S*_12_, between receive elements was measured from the preamplifier inputs.

Image SNR maps were derived from gradient-echo images, with and without RF transmission, using a covariance-weighted, root sum-of-squares reconstruction^41,58^: matrix size: 96 × 96, FOV: 240 × 240 mm, number of slices: 72, slice thickness: 2.5 mm, slice gap: 20%, TE/TR: 2.9/2,500 ms, flip angle: 7°, BW: 500 Hz/pixel, number of averages: 2. SNR maps were acquired after RF shimming over the brain with an algorithm that maximized uniformity (at the expense of efficiency) with an upper bound placed on the maximum 10-g-averaged SAR. SNR maps were then normalized by *B*_1_^+^ to remove the potential confound of transmit-related changes affecting the assessment of the receive performance.

The noise-only acquisition was additionally used to calculate the noise correlation matrix and the noise level of individual receive elements. Geometry-factor maps were created by retrospectively under-sampling the fully sampled k-space data and reconstructing using SENSE^59^. All calculations were performed in Matlab (MathWorks, Natick, MA, USA).

#### 2.3.3 Field probes

The performance of field probes was evaluated when they were located on the manufacturer’s scaffold and when integrated into the coil. When on the scaffold, field probes had a mean distance to isocentre of 9.0 cm (range: 8.2 – 9.9 cm), with their preamplifiers located approximately 70-cm away—i.e., using the manufacturer’s original cable length. *B*_0_ shimming was applied prior to all field-probe measurements: when on the scaffold, the default (“tune-up”) *B*_0_ shim was applied; when integrated into the coil, a *B*_0_ shim was calculated over the brain.

The FID amplitude and lifetime of each field probe was measured to quantify any degradation caused by integration into the coil. Since some FIDs exhibited multi-exponential decay rates, lifetimes were defined as the time required to reach 1/e (37%) of their original amplitude.

The maximum duration field probes were capable of monitoring field dynamics over was evaluated by acquiring an in-vivo, diffusion-weighted EPI sequence with a relatively long, 58-ms readout duration: FOV: 224 × 224 mm, acquisition matrix: 112 × 112, number of slices: 40, slice thickness: 2 mm, TE/TR: 124/11,000 ms, BW: 2,126 Hz/pixel, flip angle: 90°, number of averages: 3, b = 0 s/mm^2^ acquisitions: 1, diffusion directions: 16, b-value: 750 s/mm^2^, no acceleration; *B*_0_ and *B*_1_^+^ shimming was applied over the whole brain prior to the acquisition. Data from the 16 field probes was then fit with a spherical-harmonic expansion using the manufacturer’s software. Since the accuracy of the estimated spherical-harmonic terms (and the subsequent trajectory correction) was affected by the number of orders included in the fit, zeroth through first-, second-, and third-order fits were performed. The resultant field dynamics were compared to those acquired during the same pulse sequence when field probes were located on the external scaffold—all scaffold data was reconstructed using a third-order fit, as we defined this as the “gold standard” since it was the manufacturer’s suggested basis order for scans conducted with field probes on the scaffold.

An equivalent analysis was carried out for a diffusion-weighted, single-shot spiral acquisition with a four-fold acceleration rate in the anterior-posterior direction. (FOV: 192 × 192 mm, acquisition matrix: 128 × 128, number of slices: 10, slice thickness: 3 mm, TE/TR: 33/2,500 ms, BW: 2,170 Hz/pixel, flip angle: 70°, reference lines: 48, b = 0 s/mm^2^ acquisitions: 1, diffusion directions: 6, b-value: 1000 s/mm^2^.) *B*_0_ field maps were acquired for zeroth-order correction of the spiral acquisition: FOV: 240 × 240 mm, acquisition matrix: 160 × 160, number of slices: 54, slice thickness: 3 mm, TE1/TE2/TR: 4.1/5.1/1,860 ms, BW: 601 Hz/pixel, flip angle: 35°.

To assess image quality, the spiral acquisition was reconstructed with the nominal applied trajectory (i.e., “uncorrected”) and trajectories corrected for zeroth through first-, second-, and third-order field perturbations. The same acquisition was also reconstructed with zeroth through third-order field dynamics mapped while probes were located on the external scaffold.

All image reconstruction was performed in Matlab using in-house developed software^19^, with special consideration for the gradient coil, since its asymmetry produces additional *B*_0_ and linear Maxwell terms compared to symmetric coils^60^. The MRI scanner corrects for the linear terms automatically by applying dynamic gradients to counter-act them and corrects for the *B*_0_ terms by applying the appropriate demodulation phase^61^. The linear correction was presumed to be measured by the field probes, while the DC correction is invisible to the field-probe system and was thus reversed during reconstruction using Maxwell-term predictions based on the first-order field-probe measurements.

## 3 Results

The primary design objective when integrating the field probes into the coil was to minimize the impact on the performance of both the RF coil (transmit and receive) and the field probes; therefore, the performance of each subsystem was characterized independently.

### 3.1 Transmit-coil performance

The combined impedance of bazooka baluns (375 – 525 Ω) and choke baluns (> 1 kΩ) was effective in reducing common-mode currents: no sensitivity to touching or repositioning of the transmit cable was observed. Active detuning provided approximately 22 dB of isolation to the receive coil during reception.

Prior to integration of field probes, the reflection of individual elements was better than -16 dB when loaded with a head. The mean and maximum coupling between adjacent transmit elements was -28 dB and -15 dB, respectively; loops were decoupled from each other and neighbouring dipoles by -18 dB or better. After the integration of field probes, there occurred small shifts in the resonant frequency of dipoles (on the order of several MHz), which were compensated for by retuning and matching these elements. Negligible changes were observed in the resonant frequency of loop elements and inter-element coupling; therefore, no changes to capacitor values were required for the self-decoupled loops. After retuning and matching of transmit elements, the reflection of individual elements differed by one percentage point or less (when converted to a linear scale) compared to the reflection measured without field probes present. The maximum difference in coupling was less than three percentage points (Figure 3).

**Figure 3.**
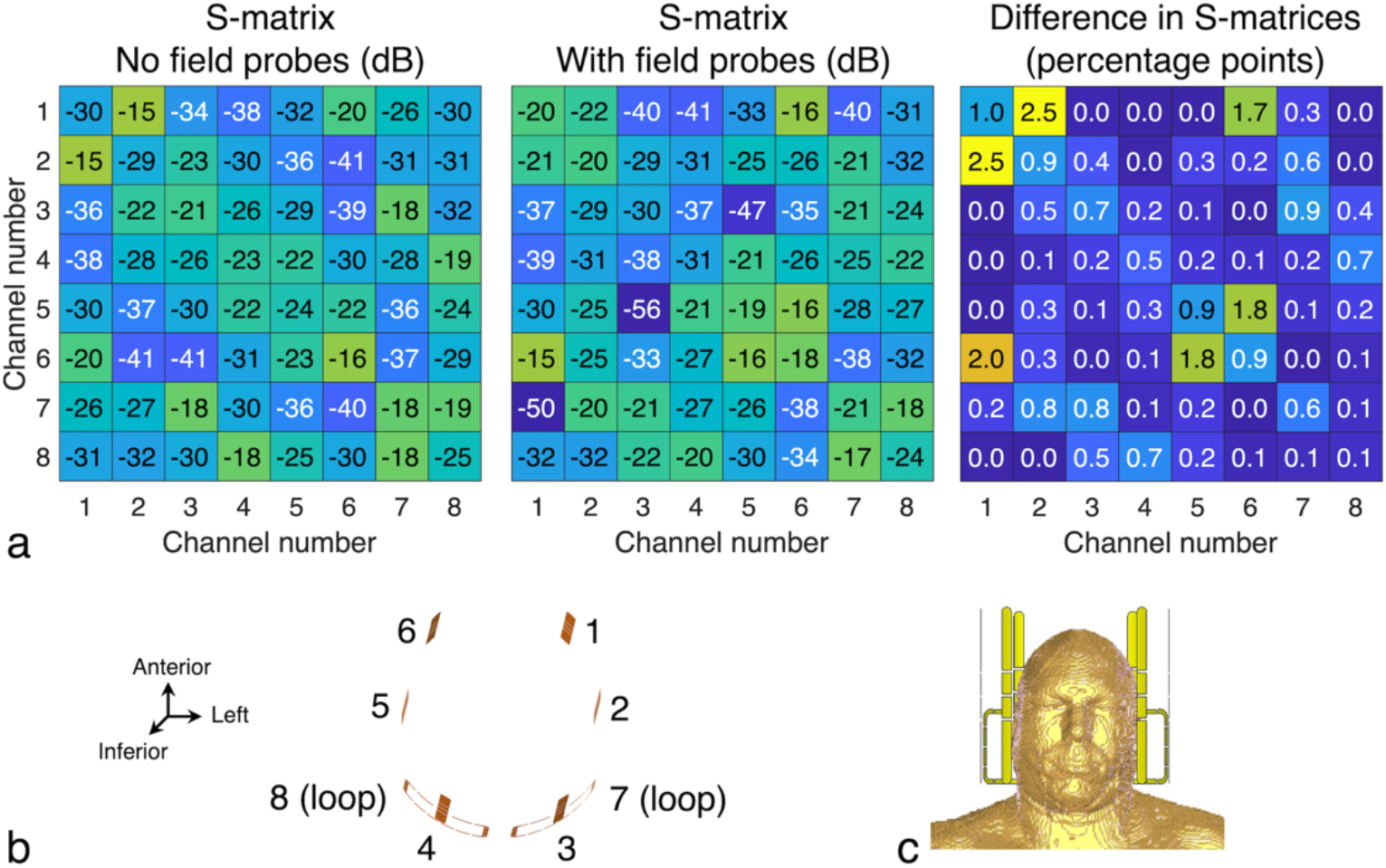
(a) The scattering-parameter matrices before and after integrating field probes into the coil, corresponding to the element positioning and numbering in (b, c). Transmit elements were retuned and matched after the integration of field probes to compensate for minor shifts in their resonant frequencies (on the order of several MHz). After converting to a linear scale, the difference in reflection was one percentage point or less and the difference in coupling was less than three percentage points.

With the integration of field probes, the *B*_1_^+^ efficiency of individual elements changed by 0.70 – 1.15-fold (when averaged over the brain), with a mean decrease over all channels of 4% (Figure 4); the spatial pattern of the *B*_1_^+^phase remained largely unchanged. RF shimming (for maximum uniformity) was capable of compensating for these small changes to the *B*_1_^+^efficiency of individual channels—the mean *B*_1_^+^efficiency of the combined map increased by 5% over the brain (from 19 nT/V to 20 nT/V). The *B*_1_^+^ uniformity over the whole brain remained constant at 26%. When taken together, this indicates the field probes had a negligible effect on the performance of the transmit coil.

**Figure 4.**
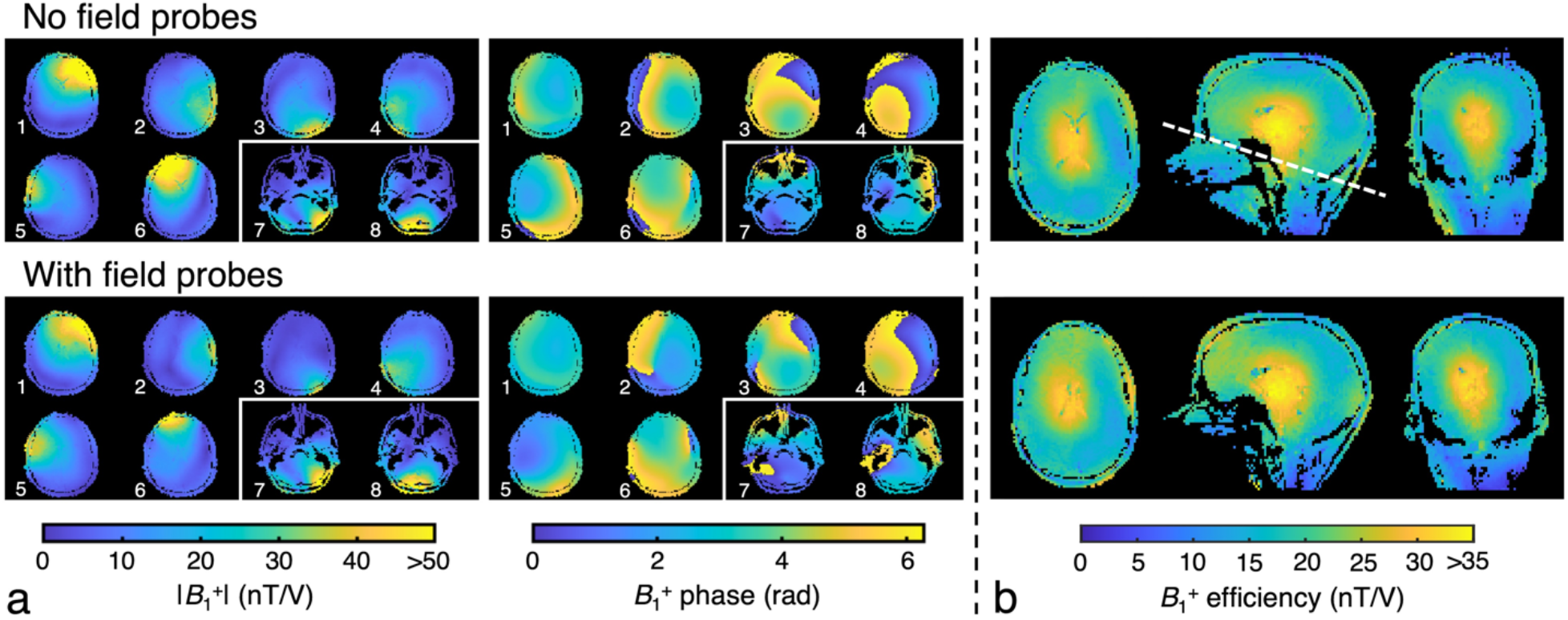
(a) Adding field probes between the transmit and receive coils produced 0.70 – 1.15-fold changes in the *B*_1_^+^ efficiency of individual elements, with a mean decrease of 4%; the spatial phase pattern of *B*_1_^+^ remained largely unaffected. (b) *B*_1_^+^ shimming was capable of compensating for small changes in the *B*_1_^+^ distribution of individual elements caused by the integration of field probes: after employing an RF shim solution that maximized uniformity—at the expense of efficiency—over the brain (in the region superior to the white dashed line), no degradation in transmit efficiency or uniformity was observed.

The *B*_1_^+^ efficiency and SAR efficiency (i.e., the *B*_1_^+^efficiency per square-root of the maximum 10-g-averaged SAR) of a maximum-efficiency shim solution is provided in Figure 5 for completeness. The *B*_1_^+^ field had a mean efficiency of 25 nT/V and a uniformity of 38% over the brain. The mean SAR efficiency was 0.38μCT/(W/kg)^1/2^, which excludes the 1.8-fold SAR safety factor implemented on the scanner.

**Figure 5.**
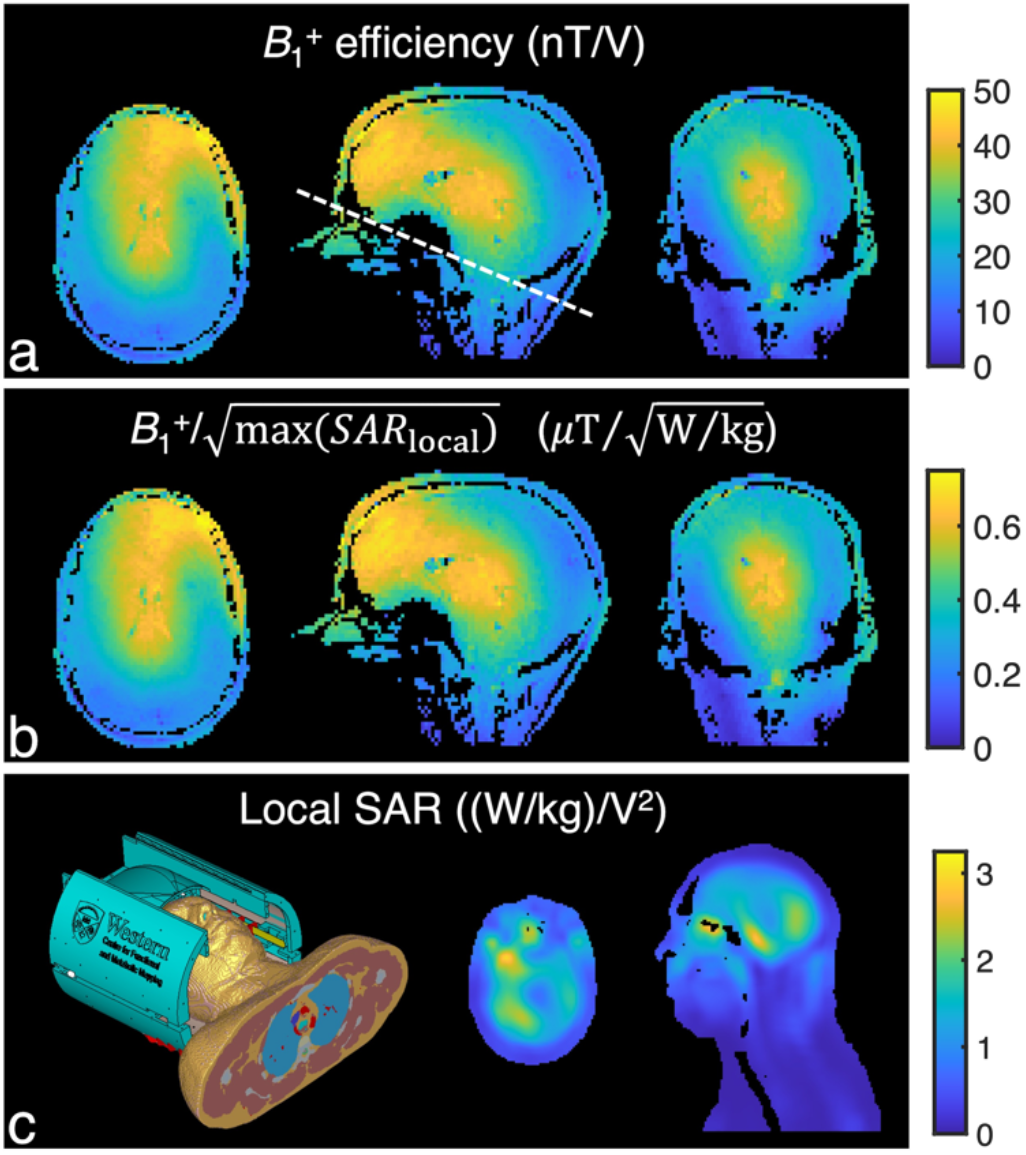
(a) In vivo *B*_1_^+^ efficiency, (b) SAR efficiency (i.e., the *B*_1_^+^efficiency per square-root of maximum 10-g-averaged SAR) resulting from an RF shim solution that maximized *B*_1_^+^ efficiency at the expense of uniformity, and (c) the 10-g-averaged SAR as calculated from the Hugo body model in CST. Performance metrics have been quoted in the text for the region superior to the white dashed line.

### 3.2 Receive-coil performance

The mean geometric decoupling between adjacent elements was -15 dB (worst-case: -10 dB). Preamplifier decoupling had a mean of 19 dB across all elements, while active detuning provided approximately 29 dB of isolation during transmission. Preamplifier decoupling and active detuning did not change after the addition of field probes. The addition of field probes did, however, cause up to a 25-Ω change in the impedance of some elements (prior to being re-tuned and matched), with affected elements residing close to the field probes.

The mean and maximum noise correlation, prior to the integration of field probes, was 11% and 53%, respectively (Figure 6a). After the integration of field probes, the mean noise correlation remained constant at 11%, while the maximum noise correlation increased by 3 percentage points. The change in normalized noise level of individual channels, before and after the integration of field probes, ranged from 0.89 – 1.14-fold, with no change in the mean taken across all channels. The difference in the mean *B*_1_^+^-corrected SNR produced by individual receive elements, before and after the integration of field probes, ranged from 0.78 – 1.15-fold with a mean change of 1% across all channels. The difference in the covariance-weighted, root sum-of-squares SNR (corrected for *B*_1_^+^) over the brain exhibited a commensurately negligible change (1%) (Figure 6b). There was additionally no change in the mean geometry factor over the brain after the addition of field probes, with a 4-fold reduction factor resulting in less than a 15% mean geometry factor (Figure 7).

**Figure 6.**
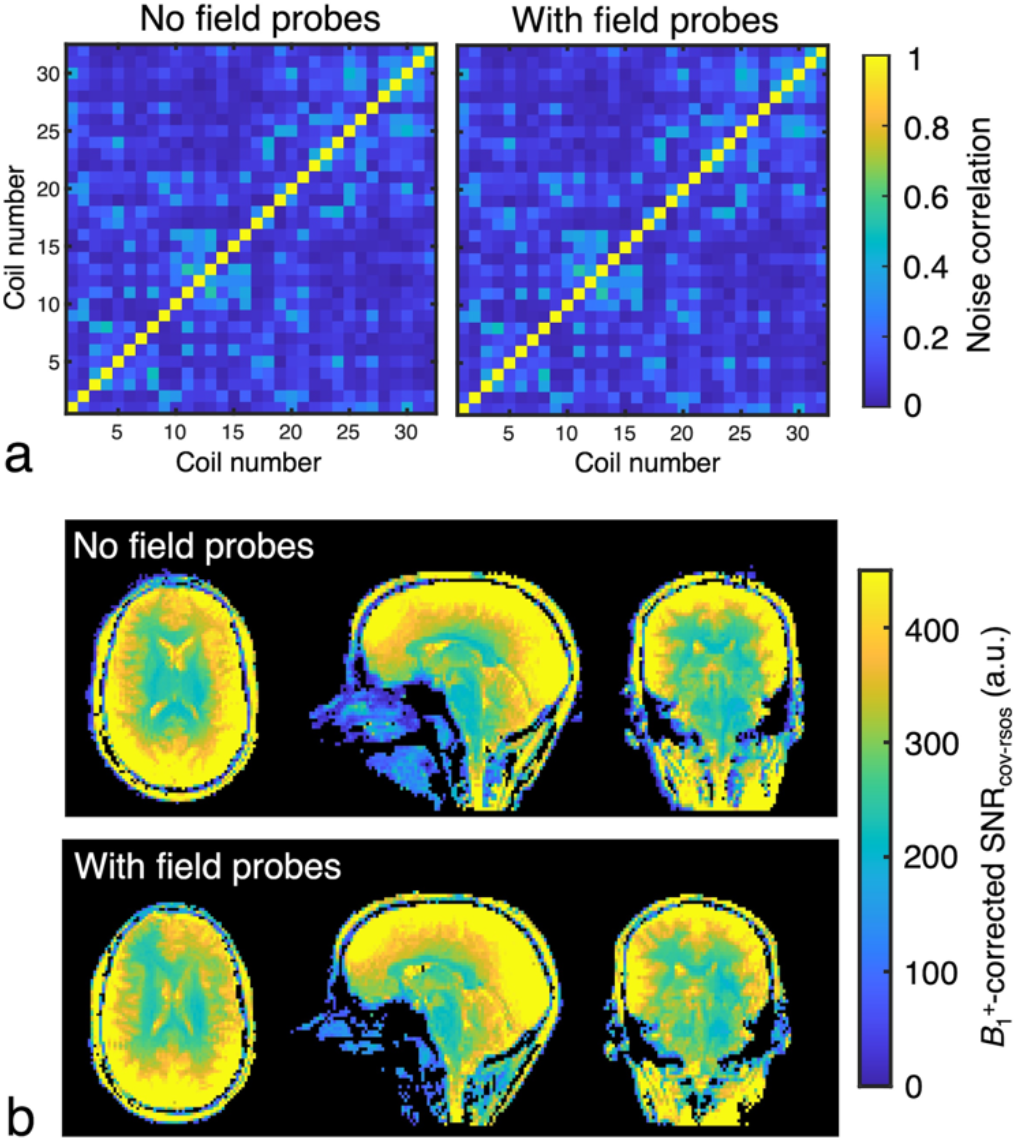
(a) Receiver noise-correlation matrices, before and after the integration of field probes into the coil, demonstrated no change in their mean values. (b) The covariance-weighted, root sum-of-squares SNR after being corrected for the spatial distribution of *B*_1_^+^. The addition of field probes into the coil caused minor differences in the spatial distribution of SNR, but no degradation of the mean SNR over the brain.

**Figure 7.**
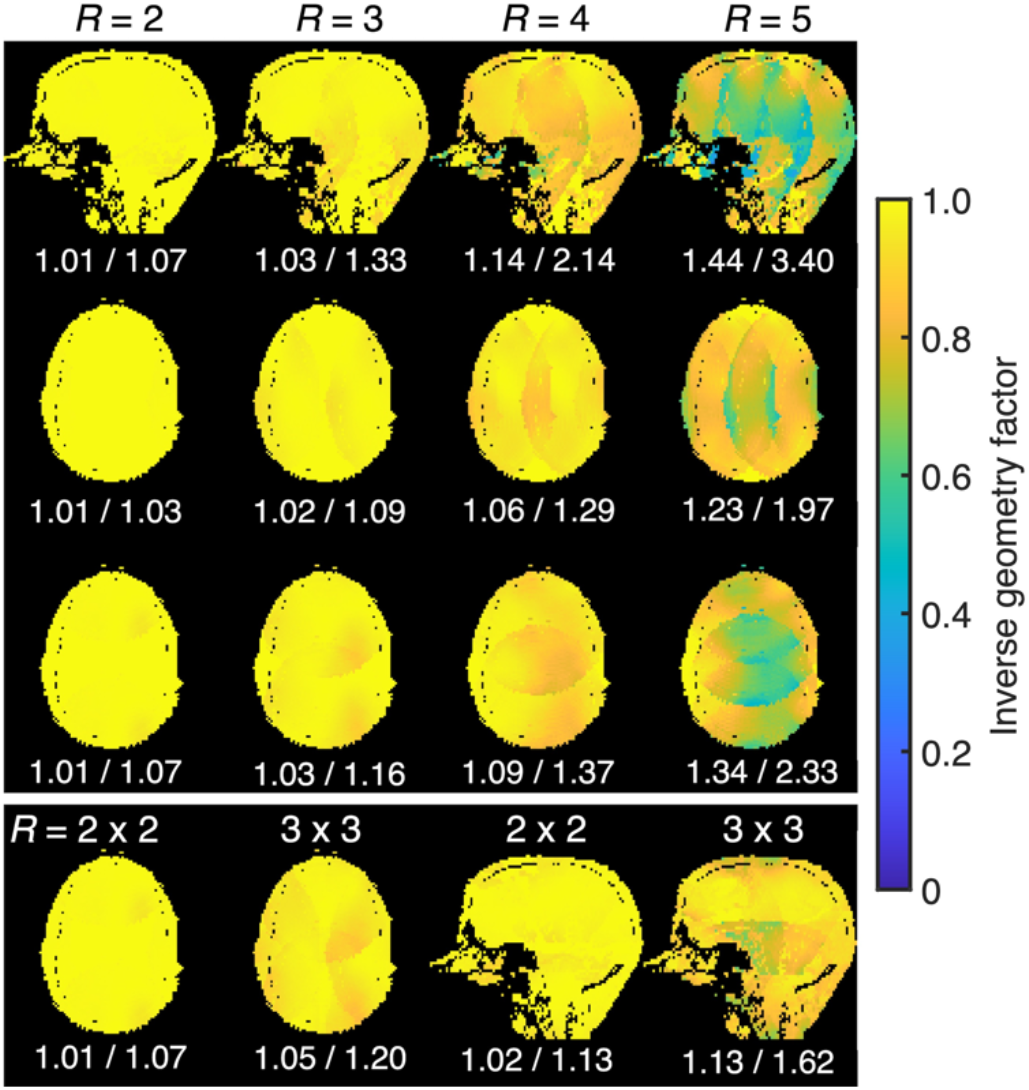
Inverse geometry factor maps acquired after the integration of field probes into the coil, with the mean and maximum geometry factor displayed beneath their respective maps. Mean geometry factors did not change with the addition of field probes into the coil. A four-fold acceleration factor could be achieved before a pronounced increase in geometry factor was observed.

### 3.3 Field-probe performance

The FID lifetimes of field probes changed by 0.55 – 1.14-fold (mean: 0.83-fold) after integration into the RF coil (Figure 8). All field probes had lifetimes (the time to reach 1/e of their original amplitude) of at least 20 ms (compared to a minimum of 34 ms when on the scaffold) and persisted for at least 40 ms before entering the noise floor of the measurement. The FID lifetimes were largely dependent on the gradient/shim currents: all probe lifetimes could be increased to near their ideal (on-scaffold) values by systematically altering the gradient/shim currents; however, ideal lifetimes could not be achieved simultaneously for all probes with a single shim setting.

**Figure 8.**
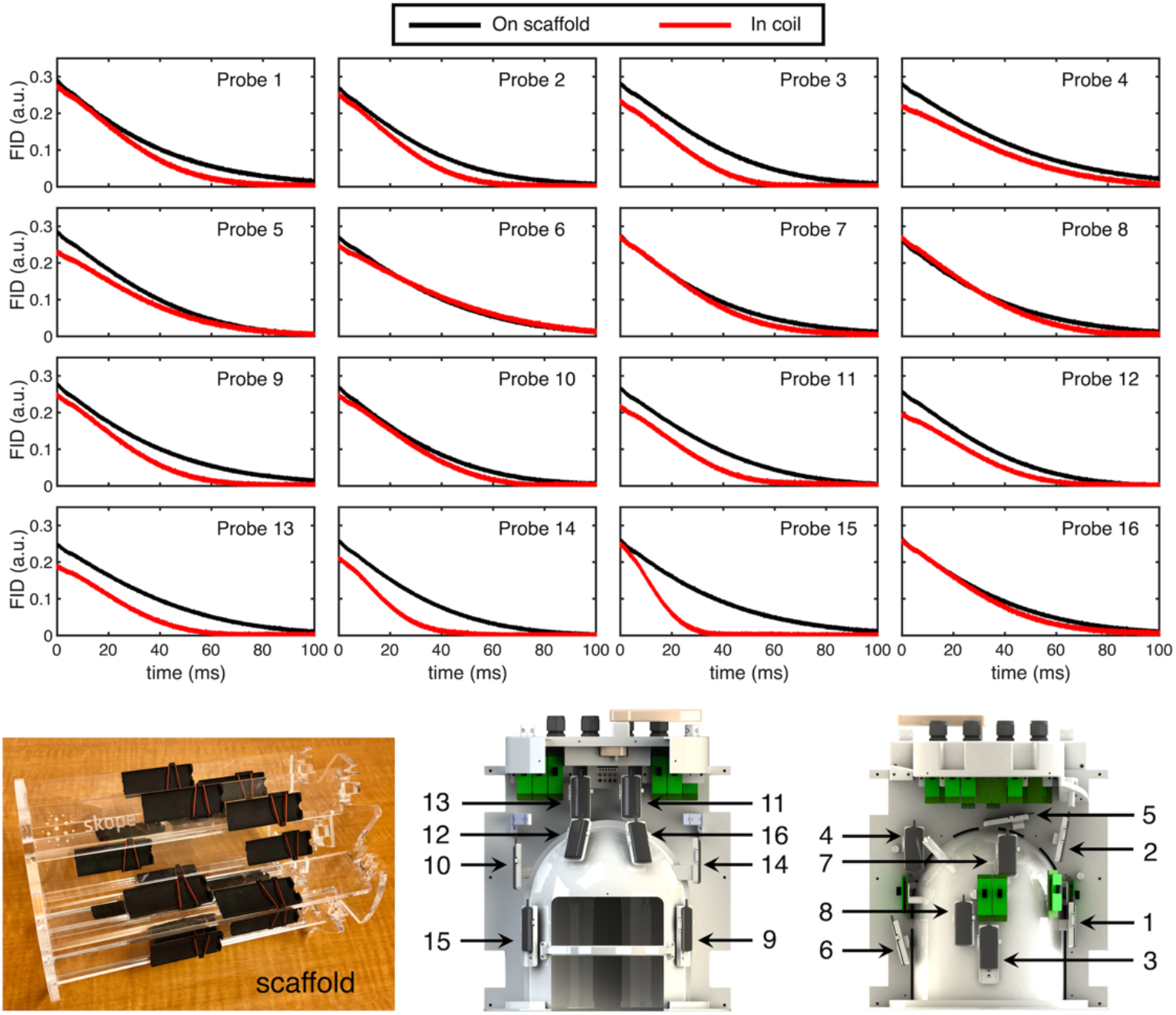
The free-induction decays of individual field probes when positioned on the manufacturer’s scaffold (i.e., in their optimal configuration) and when integrated into the coil. The applied *B*_0_ shim had the greatest influence on FID lifetime—a consequence of the reduced region of gradient- and shim-field fidelity within a head-only gradient coil. Field-probe numbering corresponds to positions provided in the CAD drawing (lower right).

The FID amplitudes changed by 0.77 – 1.06-fold (mean: 0.90-fold), indicating the additional cabling required to place the field-probe preamplifiers outside of the scanner resulted in a modest, yet tolerable, loss of signal.

The field probe with the shortest lifetime dictated the duration in k-space that could be measured with precision—approximately 40 – 45 ms (Figures 9a, b). This allowed field dynamics to be measured for at least 40 ms (Figure 9c, d) during an EPI acquisition, which was considerably longer than the 13-ms readout duration of spiral acquisitions acquired in this study. When field probes were incorporated into the coil, a systematic error was introduced into the determination of spherical-harmonic terms (compared to scaffold data) when employing a third-order fit to the field-probe data (Figure 9d); this error was greatly reduced when limiting the fit to first or second order. An equivalent analysis for the shorter spiral acquisition demonstrated the same trend, albeit with a lower error with respect to scaffold data due to the shorter acquisition time (see Supporting Information Figure S1).

**Figure 9.**
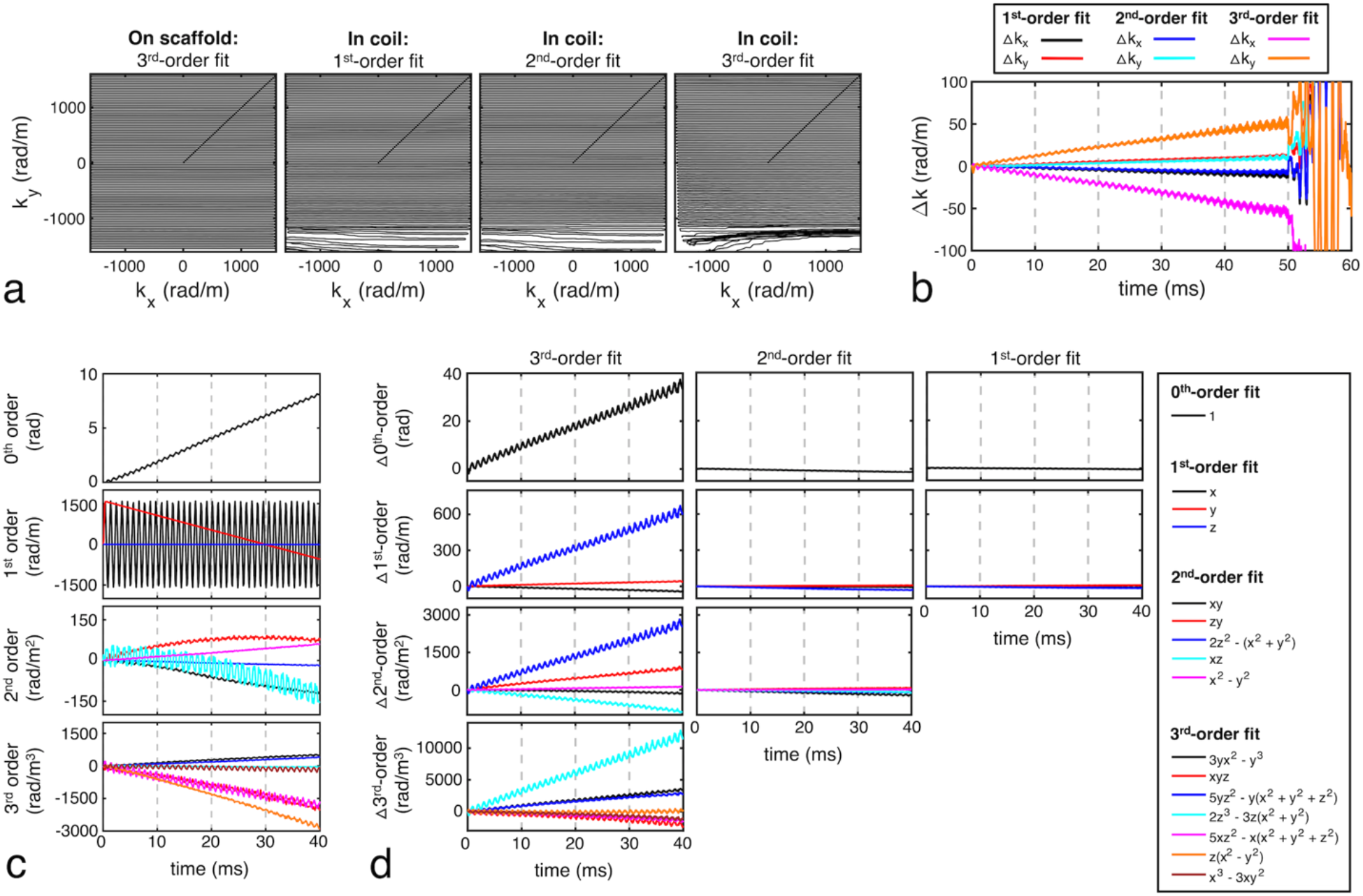
(a) K-space of a diffusion-weighted EPI acquisition as measured by field probes when on the scaffold and when integrated into the coil (for first-, second-, and third-order spherical-harmonic fits to the field-probe data) and (b) their commensurate difference (Δk = k_scaffold_ – k_coil_). (c) Field dynamics measured by probes while on the scaffold, and corrected with a third-order fit, were used as a reference. (d) The difference between these field dynamics and those measured while field probes were integrated into the coil exhibited a systematic deviation when employing a third-order fit: this was most prominent for terms with a strong *z*-dependency, most likely due to the asymmetry of the gradient coil. These errors were substantially reduced when employing a first- or second-order fit. Integrated field probes were capable of measuring field dynamics for a minimum duration of 40 ms, which is commensurate with the shortest FID duration (see Figure 8).

Spiral images reconstructed with trajectories measured with integrated probes and corrected for zeroth through first- and second-order field perturbations demonstrated improvements in image quality (reduced blurring) when compared to uncorrected images (Figure 10). When employing a diffusion-weighted reconstruction with up to first-order correction of field perturbations, images were qualitatively similar to those corrected with a third-order fit to scaffold data; non-diffusion-weighted reconstructions achieved similar image quality up to a second-order fit. Relative differences between images corrected with integrated versus scaffold-mounted field probes manifested primarily at tissue boundaries. These relative differences were exacerbated when employing a third-order correction with integrated probes, which is congruent with the larger error in measured field dynamics for third-order fits (Supporting Information Figure S1d).

**Figure 10.**
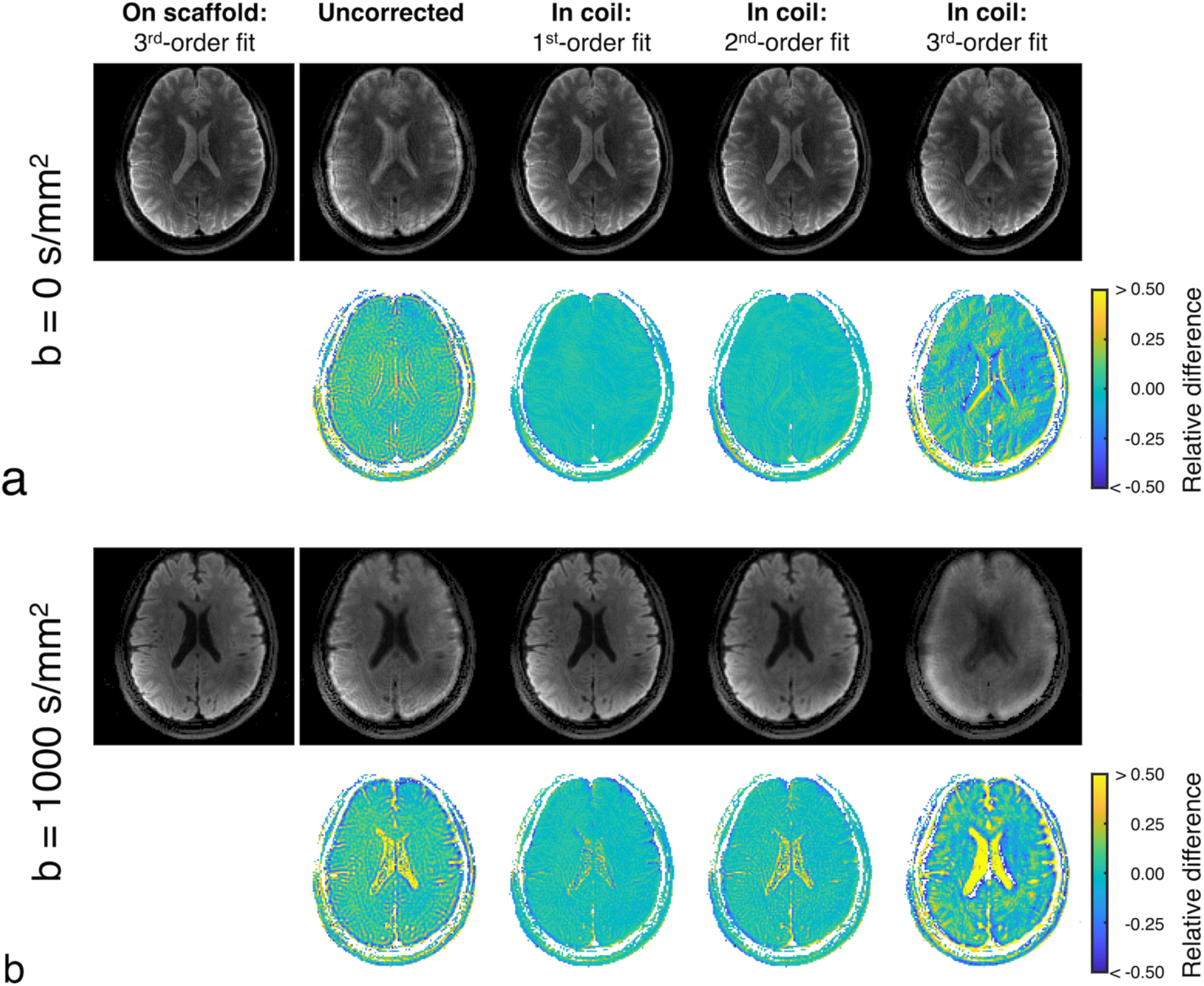
A spiral acquisition with (a) non-diffusion-weighted (b = 0 s/mm^2^) and (b) diffusion-weighted (b = 1000 s/mm^2^) reconstructions. Images are shown for an uncorrected trajectory as well as for trajectories corrected for zeroth through first-, second-, and third-order field perturbations, as measured with integrated field probes. As a reference, the same image was reconstructed with trajectories measured by field probes mounted on the external scaffold and corrected for zeroth through third-order field perturbations—this image was used as the baseline to calculate the relative difference to all other image reconstructions.

## 4 Discussion

This study presents a solution for integrating a commercial field camera into an RF coil for ultra-high field imaging. This permits the characterization and correction of field perturbations that can otherwise reduce the fidelity of k-space trajectories and degrade image quality. Diffusion-weighted imaging, a gradient-sensitive sequence, is particularly amenable to this technique and further benefits from ultra-high field strengths and head-only gradient coils. These technological advancements nonetheless impose technical challenges on the development of an RF coil with an integrated field camera—a limited bore space and a reduced region of gradient- and shim-field fidelity. By designing the RF coil with the original intent to include field probes, these challenges could be considered during the design stage to improve performance.

### 4.1 RF-coil performance

Careful planning of field-probe positions and cabling were undertaken to tailor their placement to the head while also avoiding proximity to regions potentially detrimental to the performance of the RF coil (directly beneath transmit elements; near receive cabling; above the matching, active-detuning, and passive-detuning circuits of receive elements). These design considerations resulted in the field probes being almost entirely transparent to the transmit and receive coils, despite their proximity necessitated by their use in a head-only gradient coil. No loss in transmit performance (efficiency and uniformity) or receive performance (noise correlation, SNR, geometry factor) was observed.

The transmit coil produced a *B*_1_^+^ field with sufficient uniformity for diffusion-weighted imaging—a feat made difficult by high flip-angle excitation and refocusing pulses. The geometrical layout, with two loops near the cerebellum, increased *B*_1_^+^ efficiency in this region and permitted an additional degree of freedom when RF shimming along the superior-inferior direction.

The receive-coil layout permitted realistic acceleration rates up to four-fold before excessive noise amplification (i.e., high geometry factors) occurred during reconstruction. This allowed the shortening of echo trains to allow the readout duration to be less than the lifetime of field probes.

### 4.2 Field-camera performance

In this study, the reduced imaging region of the head-only gradient coil had the greatest impact on the efficacy of the field camera in characterizing higher-order field dynamics. The positioning of field probes in the non-linear region of the gradient coil resulted in inaccuracies in the measured probe positions (0.3 – 1.8 cm) and spatially non-linear phase accrual of FID phases. The combination of these inaccuracies resulted in third-order fits becoming erroneous, as the 16 field probes could no longer adequately characterize all 16 spherical-harmonic field terms. First- and second-order fits, however, demonstrated a considerably lower deviation from scaffold data (Figure 9d). The low relative difference between images corrected with first- and second-order fits using integrated field probes and third-order fits using scaffold-mounted field probes suggests diminishing benefits of higher-order field corrections in a well-optimized scanner (Figure 10). However, the improvement realized with higher-order field corrections could likely be increased by incorporating the spherical-harmonic representation of non-linearity of the gradient field into the correction algorithm—provided that the magnetic field varies monotonically at the location of the field probes—thereby compensating for the aforementioned ramifications of placing the field probes in the non-linear region of the head-only gradient coil.

Overall, by integrating the field camera into the coil, scanning workflow is greatly improved and the design architecture now facilitates subject-specific field corrections caused by motion— which can be more prominent for longer acquisition protocols^33^.

## Acknowledgements

The authors would like to thank Paul Weavers and Cameron Cushing from Skope for helpful discussions and Alexander Mashkovtsev and Peter Zeman for fabrication assistance. This work was supported by the Canada Foundation for Innovation (to RSM); Canada First Research Excellence Fund to BrainsCAN; Brain Canada Platform Support Grant (to RSM); Natural Sciences and Engineering Research Council of Canada Discovery Grants (to CAB [RGPIN-2018-05448] and RSM [RGPIN-2019-04743]) and the Robarts Research Institute.

## Data availability statement

The data that support the findings of this study are openly available in an OSF repository (DOI: 10.17605/OSF.IO/7TSJM). This includes CAD files of the RF coil and all performance and imaging data.

---

**Supporting Information Figure S1.**
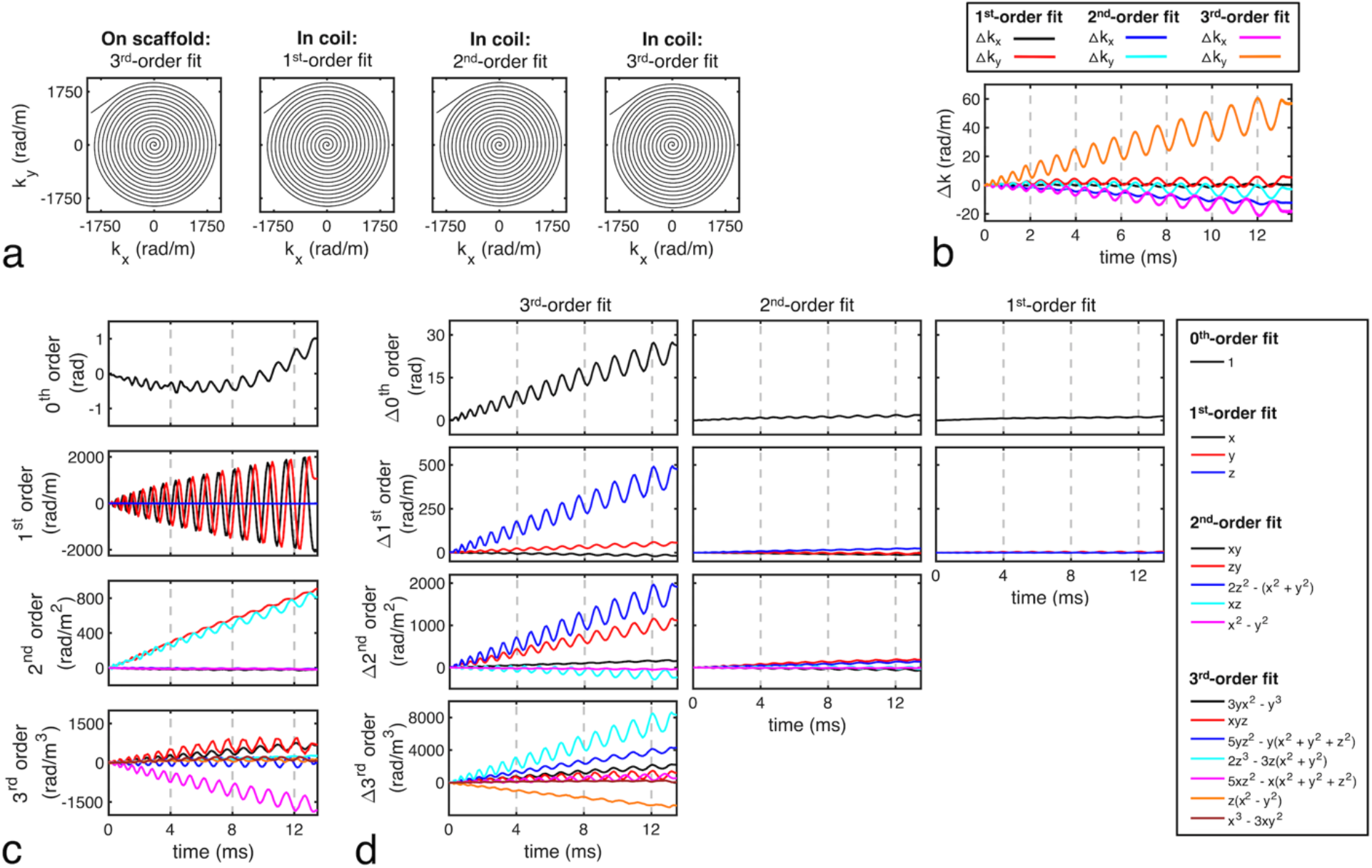
(a) K-space of a diffusion-weighted, single-shot spiral acquisition as measured by field probes when on the scaffold and when integrated into the coil (for first-, second-, and third-order spherical-harmonic fits to the field-probe data) and (b) their commensurate difference (Δk = k_scaffold_ – k_coil_). (c) Field dynamics measured by probes while on the scaffold, and corrected with a third-order fit, were used as a reference. (d) The difference between these field dynamics and those measured while field probes were integrated into the coil exhibited a systematic deviation when employing a third-order fit: this was most prominent for terms with a strong *z*-dependency, most likely due to the asymmetry of the gradient coil. These errors were substantially reduced when employing a first- or second-order fit.

## References

1. Barmet C, De Zanche N, Pruessmann KP. Spatiotemporal magnetic field monitoring for MR. Magn Reson Med. 2008;60(1):187–197.

2. Boesch C, Gruetter R, Martin E. Temporal and spatial analysis of fields generated by eddy currents in superconducting magnets: optimization of corrections and quantitative characterization of magnet/gradient systems. Magn Reson Med. 1991;20(2):268–284.

3. Morich MA, Lampman DA, Dannels WR, Goldie FD. Exact temporal eddy current compensation in magnetic resonance imaging systems. IEEE Trans Med Imaging. 1988;7(3):247–254.

4. Ebel A, Maudsley AA. Detection and correction of frequency instabilities for volumetric 1H echo-planar spectroscopic imaging. Magn Reson Med. 2005;53(2):465–469.

5. Wu Y, Chronik BA, Bowen C, Mechefske CK, Rutt BK. Gradient-induced acoustic and magnetic field fluctuations in a 4T whole-body MR imager. Magn Reson Med. 2000;44(4):532–536.

6. Zaitsev M, Maclaren J, Herbst M. Motion artifacts in MRI: A complex problem with many partial solutions. J Magn Reson Imaging. 2015;42(4):887–901.

7. Duyn JH, Yang Y, Frank JA, van der Veen JW. Simple correction method for k-space trajectory deviations in MRI. J Magn Reson. 1998;132(1):150–153.

8. Giese D, Haeberlin M, Barmet C, Pruessmann KP, Schaeffter T, Kozerke S. Analysis and correction of background velocity offsets in phase-contrast flow measurements using magnetic field monitoring. Magn Reson Med. 2012;67(5):1294–1302.

9. Jang H, McMillan AB. A rapid and robust gradient measurement technique using dynamic single-point imaging. Magn Reson Med. 2017;78(3):950–962.

10. De Zanche N, Barmet C, Nordmeyer-Massner JA, Pruessmann KP. NMR probes for measuring magnetic fields and field dynamics in MR systems. Magn Reson Med. 2008;60(1):176–186.

11. Vannesjo SJ, Wilm BJ, Duerst Y, et al. Retrospective correction of physiological field fluctuations in high-field brain MRI using concurrent field monitoring. Magn Reson Med. 2015;73(5):1833–1843.

12. Barmet C, De Zanche N, Wilm BJ, Pruessmann KP. A transmit/receive system for magnetic field monitoring of in vivo MRI. Magn Reson Med. 2009;62(1):269–276.

13. Sipilä P, Greding S, Wachutka G, Wiesinger F. 2H transmit-receive NMR probes for magnetic field monitoring in MRI. Magn Reson Med. 2011;65(5):1498–1506.

14. Sipilä P, Lange D, Lechner S, et al. Robust, susceptibility-matched NMR probes for compensation of magnetic field imperfections in magnetic resonance imaging (MRI). Sens Actuators A Phys. 2008;145-146:139–146.

15. Handwerker J, Bonehi V, Eschelbach M, Scheffler K, Ortmanns M, Anders J. An Active Transmit/Receive NMR Magnetometer for Field Monitoring in Ultra High Field MRI Scanners. Biomed Tech. 2013;58 Suppl 1. doi:10.1515/bmt-2013-4263

16. Wehkamp N, Rovedo P, Fischer E, Hennig J, Zaitsev M. Frequency-adjustable magnetic field probes. Magn Reson Med. 2021;85(2):1123–1133.

17. Boero G, Frounchi J, Furrer B, Besse P-A, Popovic RS. Fully integrated probe for proton nuclear magnetic resonance magnetometry. Rev Sci Instrum. 2001;72(6):2764–2768.

18. Dietrich BE, Brunner DO, Wilm BJ, et al. A field camera for MR sequence monitoring and system analysis. Magn Reson Med. 2016;75(4):1831–1840.

19. Wilm BJ, Barmet C, Pavan M, Pruessmann KP. Higher order reconstruction for MRI in the presence of spatiotemporal field perturbations. Magn Reson Med. 2011;65(6):1690–1701.

20. Barmet C, Wilm BJ, Pavan M, Pruessmann KP. A third-order field camera with microsecond resolution for MR system diagnostics. In: Proceedings of the 17th Annual Meeting of ISMRM. Honolulu, USA; 2009:781.

21. Vannesjo SJ, Haeberlin M, Kasper L, et al. Gradient system characterization by impulse response measurements with a dynamic field camera. Magn Reson Med. 2013;69(2):583–593.

22. Ma R, Akçakaya M, Moeller S, Auerbach E, Uğurbil K, Van de Moortele P-F. A field-monitoring-based approach for correcting eddy-current-induced artifacts of up to the 2nd spatial order in human-connectome-project-style multiband diffusion MRI experiment at 7T: A pilot study. Neuroimage. 2020;216:116861.

23. Stich M, Pfaff C, Wech T, et al. The temperature dependence of gradient system response characteristics. Magn Reson Med. 2020;83(4):1519–1527.

24. Brunheim S, Mirkes C, Dietrich BE, et al. Replaceable field probe holder for the Nova coil on a 7 Tesla Siemens scanner. In: Proceedings of the 28th Annual Meeting of ISMRM.; 2020:3389.

25. Kennedy M, Lee Y, Nagy Z. An industrial design solution for integrating NMR magnetic field sensors into an MRI scanner. Magn Reson Med. 2018;80(2):833–839.

26. Mahmutovic M, Scholz A, Kutscha N, et al. A 64-Channel Brain Array Coil with an Integrated 16-Channel Field Monitoring System for 3T MRI. In: Proceedings of the 29th Annual Meeting of ISMRM. ; 2021:0623.

27. Brunner DO, Gross S, Schmid T, et al. Integration of field monitoring for neuroscientific applications - SNR, acceleration and image integrity. In: Proceedings of the 27th Annual Meeting of ISMRM. Montreal, Canada; 2019:1046.

28. Bollmann S, Kasper L, Vannesjo SJ, et al. Analysis and correction of field fluctuations in fMRI data using field monitoring. Neuroimage. 2017;154:92–105.

29. Wilm BJ, Barmet C, Gross S, et al. Single-shot spiral imaging enabled by an expanded encoding model: Demonstration in diffusion MRI. Magn Reson Med. 2017;77(1):83–91.

30. Kasper L, Engel M, Barmet C, et al. Rapid anatomical brain imaging using spiral acquisition and an expanded signal model. Neuroimage. 2018;168:88–100.

31. Le Bihan D. Looking into the functional architecture of the brain with diffusion MRI. Nat Rev Neurosci. 2003;4(6):469–480.

32. Le Bihan D, Poupon C, Amadon A, Lethimonnier F. Artifacts and pitfalls in diffusion MRI. J Magn Reson Imaging. 2006;24(3):478–488.

33. Lee Y, Wilm BJ, Brunner DO, et al. On the signal-to-noise ratio benefit of spiral acquisition in diffusion MRI. Magn Reson Med. 2021;85(4):1924–1937.

34. Wilm BJ, Nagy Z, Barmet C, et al. Diffusion MRI with concurrent magnetic field monitoring. Magn Reson Med. 2015;74(4):925–933.

35. Sotiropoulos SN, Moeller S, Jbabdi S, et al. Effects of image reconstruction on fiber orientation mapping from multichannel diffusion MRI: reducing the noise floor using SENSE. Magn Reson Med. 2013;70(6):1682–1689.

36. Foo TKF, Tan ET, Vermilyea ME, et al. Highly efficient head-only magnetic field insert gradient coil for achieving simultaneous high gradient amplitude and slew rate at 3.0T (MAGNUS) for brain microstructure imaging. Magn Reson Med. 2020;83(6):2356–2369.

37. Setsompop K, Kimmlingen R, Eberlein E, et al. Pushing the limits of in vivo diffusion MRI for the Human Connectome Project. Neuroimage. 2013;80:220–233.

38. McNab JA, Edlow BL, Witzel T, et al. The Human Connectome Project and beyond: initial applications of 300 mT/m gradients. Neuroimage. 2013;80:234–245.

39. Feinberg DA, Dietz P, Liu C, et al. Design and Development of a Next-Generation 7T human brain scanner with high-performance gradient coil and dense RF arrays. In: Proceedings of the 29th Annual Meeting of ISMRM. ; 2021:0562.

40. Gilbert KM, Klassen LM, Mashkovtsev A, Zeman P, Menon RS, Gati JS. Radiofrequency coil for routine ultra-high-field imaging with an unobstructed visual field. NMR Biomed. 2021;34(3):e4457.

41. Roemer PB, Edelstein WA, Hayes CE, Souza SP, Mueller OM. The NMR phased array. Magn Reson Med. 1990;16(2):192–225.

42. Keil B, Blau JN, Biber S, et al. A 64-channel 3T array coil for accelerated brain MRI. Magn Reson Med. 2013;70(1):248–258.

43. Gilbert KM, Curtis AT, Gati JS, Klassen LM, Menon RS. A radiofrequency coil to facilitate B1+ shimming and parallel imaging acceleration in three dimensions at 7 T. NMR Biomed. 2011;24(7):815–823.

44. Gilbert KM, Belliveau J-G, Curtis AT, Gati JS, Klassen LM, Menon RS. A conformal transceive array for 7 T neuroimaging. Magn Reson Med. 2012;67(5):1487–1496.

45. Yan X, Gore JC, Grissom WA. Self-decoupled radiofrequency coils for magnetic resonance imaging. Nat Commun. 2018;9(1):3481.

46. Eichfelder G, Gebhardt M. Local specific absorption rate control for parallel transmission by virtual observation points. Magn Reson Med. 2011;66(5):1468–1476.

47. Wiggins GC, Triantafyllou C, Potthast A, Reykowski A, Nittka M, Wald LL. 32-channel 3 Tesla receive-only phased-array head coil with soccer-ball element geometry. Magn Reson Med. 2006;56(1):216–223.

48. Burl M, Zou MX. Transmit mode coil detuning for MRI systems. US Patent 6,850,067. February 2005.

49. Chu Y-H, Hsu Y-C, Lin F-H. Decoupled dynamic magnetic field measurements improves diffusion-weighted magnetic resonance images. Sci Rep. 2017;7(1):11630.

50. Seeber DA, Jevtic J, Menon A. Floating shield current suppression trap. Concepts Magn Reson. 2004;21B(1):26–31.

51. International Electrotechnical Commission. Medical Electrical Equipment – Part 1: General Requirements for Basic Safety and Essential Performance Amendment 1. Geneva: International Electrotechnical Commission; 2012.

52. International Electrotechnical Commission. Medical Electrical Equipment – Part 2-33: Particular Requirements for the Basic Safety and Essential Performance of Magnetic Resonance Equipment for Medical Diagnosis, Edition 3.1. Geneva: International Electrotechnical Commission; 2013.

53. Hoffmann J, Henning A, Giapitzakis IA, et al. Safety testing and operational procedures for self-developed radiofrequency coils. NMR Biomed. 2016;29(9):1131–1144.

54. Yarnykh VL. Actual flip-angle imaging in the pulsed steady state: a method for rapid three-dimensional mapping of the transmitted radiofrequency field. Magn Reson Med. 2007;57(1):192–200.

55. Nehrke K. On the steady-state properties of actual flip angle imaging (AFI). Magn Reson Med. 2009;61(1):84–92.

56. Duan Q, Duyn JH, Gudino N, et al. Characterization of a dielectric phantom for high-field magnetic resonance imaging applications. Med Phys. 2014;41(10):102303.

57. Keil B, Alagappan V, Mareyam A, et al. Size-optimized 32-channel brain arrays for 3 T pediatric imaging. Magn Reson Med. 2011;66(6):1777–1787.

58. Kellman P, McVeigh ER. Image reconstruction in SNR units: a general method for SNR measurement. Magn Reson Med. 2005;54(6):1439–1447.

59. Pruessmann KP, Weiger M, Scheidegger MB, Boesiger P. SENSE: sensitivity encoding for fast MRI. Magn Reson Med. 1999;42(5):952–962.

60. Meier C, Zwanger M, Feiweier T, Porter D. Concomitant field terms for asymmetric gradient coils: consequences for diffusion, flow, and echo-planar imaging. Magn Reson Med. 2008;60(1):128–134.

61. Weavers PT, Tao S, Trzasko JD, et al. B0 concomitant field compensation for MRI systems employing asymmetric transverse gradient coils. Magn Reson Med. 2018;79(3):1538–1544.

